# Comparative benchmarking of single-cell transcriptomes and immune repertoires across technologies

**DOI:** 10.64898/2026.04.20.719117

**Authors:** Cathal King, Munir Iqbal, Elham Shokati, Celine Man Ying Li, Runhao Li, Yoko Tomita, Eric Smith, Joanna Achinger-Kawecka, Sen Wang, Kevin Fenix

## Abstract

Immune receptor profiling enables tracking of individual T or B cell clones across time and tissues, providing insight into immune responses, cancer, and autoimmunity. When combined with single-cell transcriptomics, it links clonotype identity to cellular function, revealing the diversity and dynamics of immune cell populations. This study presents a head-to-head benchmarking comparison of two single-cell immune profiling technologies: droplet-based microfluidics from 10x Genomics (10x) and combinatorial barcoding from Parse Biosciences (Parse). Using matched human samples from PBMC’s, the analysis evaluates performance across transcriptomic and T cell immune receptor features to assess data quality, reproducibility, and chain-specific recovery. The findings provide a framework for interpreting single-cell immune profiling platforms and emphasize the importance of accounting for technology-specific biases in bioinformatic analyses.

## 1. Introduction

Single-cell RNA sequencing (scRNA-seq) has become a cornerstone of modern genomics, enabling high-resolution analysis of cellular heterogeneity across complex tissues and systems. Recent technological advances now allow the profiling of millions of cells per experiment with improved sensitivity, yield, and workflow efficiency. These developments have also reduced sequencing costs, further accelerated by alternative platforms such as MGI and Ultima, making large-scale single-cell studies increasingly accessible.

The main technologies used in scRNA-seq today can be broadly classified into four categories: (1) emulsion-based methods, where cells are encapsulated into droplets; (2) microwell-based methods, where cells are physically separated into wells that serve as reaction chambers; (3) combinatorial barcoding methods, which use fixed cells as reaction chambers and sequentially add unique barcodes without physical isolation; and (4) hydrogel-based methods, which embed cells with barcoded capture beads within a hydrogel matrix. These approaches vary in throughput and sensitivity, which influences data interpretation and cross-platform comparisons.

Single-cell immune profiling captures transcriptome-wide mRNA expression (GEX) and full-length TCR or BCR sequences, enabling characterization of immune receptor diversity at single-cell resolution. Immune receptor profiling using single-cell sequencing captures paired T-cell receptor (TCR) or B-cell receptor (BCR) sequences alongside transcriptomes, linking the immune repertoire to cell type and gene expression. This enables detailed analysis of clonal expansion, immune repertoire diversity, and antigen-driven responses in cancer, infection, and autoimmunity, while connecting clonotype identity with transcriptional clustering.

T cell antigen specificity and clonal identity are defined by the TCR, which exists as either an α/β or γ/δ heterodimer, with α/β TCRs predominating on CD4_+_ helper and CD8CD4_+_ cytotoxic T cells. The extensive diversity of the TCR arises through somatic recombination of variable (V), diversity (D), and joining (J) gene segments, which together determine the sequence of the Complementarity-Determining Region 3 (CDR3)—the primary region responsible for antigen recognition in the context of the Major Histocompatibility Complex (MHC) [1]. In humans, the TCRα locus comprises 50 functional Vα and 61 Jα segments, while the TCRβ locus includes 52 Vβ, 2 Dβ, and 13 Jβ segments. Additional diversity is generated through random nucleotide insertions and deletions at V(D)J junctions, resulting in an estimated repertoire of 10^1^ –10^2^ possible TCR α/β combinations [3].

Selecting the most effective technology and strategy is essential to ensure that resulting data are both high-quality and biologically meaningful. Practical factors such as cost, hands-on time, protocol complexity, and sample compatibility also shape platform choice. Bench-marking across technologies is therefore critical to identify technical biases, improve reproducibility, and support accurate comparison and integration of single-cell immune profiling datasets.

## 2. Methods

### 2.1. Study design

Here, we performed benchmarking of two commonly used single-cell TCR kits. The kits used for this analysis were Parse Evercode™ Paired TCR + WT (v3) and 10x Genomics Chromium Next GEM Single Cell 5’ V(D)J (v2). To evaluate the performance of both immune profiling kits, we conducted a benchmark study using matched biological and technical replicates of cytokine induced killer (CIK) cells expanded from peripheral blood mononuclear cells (PBMCs) isolated from treatment-naive metastatic colorectal cancer (CRC) patients. All samples represent day 14 time points of *in vitro*-expanded CIK cells using a previously established CIK cells expansion protocol [2]. The Human Research Ethics Committee of the Central Adelaide Local Health Network granted ethical approval for this research. (HREC/14/TQEHLMH/164).

**Table 1.** Matched samples between Parse Evercode and 10x Genomics.

| Sample ID | Parse Evercode | 10x Genomics |
| --- | --- | --- |
| 234 | Parse_234 | 10X_234 |
| 236 | Parse_236 | 10X_236 |
| 237 | Parse_237 | 10X_237 |
| 238 | Parse_238 | 10X_238 |

### 2.2. Sample preparation

All samples were prepared according to the manufacturers’ protocols. For Parse samples, cells were fixed using the Evercode Cell Fixation v3 protocol following CIK cell generation on day 14 and stored at −80°C until all four samples were collected. Library preparation was then performed simultaneously for all fixed samples prior to sequencing (MGI DNBSEQ-T7). In contrast, for 10x, day 14 CIK cells were collected and freshly sent for library preparation. Libraries were generated individually per sample following manufacturer protocols, then pooled prior to paired-end sequencing (MGI DNBSEQ-G400).

### 2.3. Data processing and Analyses

Data processing pipelines and software provided by each vendor was used to pre-process, align and produce gene by cell count matrices and immune repertoire data. Bioinformatic pipelines were run with default parameters unless specified.

For Parse data, `split-pipe` version 1.4.1 was used. As this data was generated together with a whole transcriptome (WT) parent, the results from the WT data is processed first and then used for filtering for high confidence cells in the subsequent TCR analysis. This results in a more accurate representation of the TCR repertoire than processing TCR data without a parent. Therefore, WT data was first processed from each sub-library individually with the step `mode-all`. Then, the outputs from each sub-library are combined with `mode-combine`. TCR FASTQ’s were then processed by sub-library alongside each WT parent.

For 10x data, `Cell Ranger` version 9.0.1 was used. The sub-command `multi` was used which analyses multiple library types together and is the recommend pipeline for processing 5’ data. `Cell Ranger multi`, in a similar fashion to `split-pipe` for Parse data, enables more consistent cell calling by using cell calls provided by the gene expression data to improve cell calls from the V(D)J data.

Various frameworks are now available for downstream analysis of single-cell data. These tools are primarily built in R or Python, with examples including Seurat [4] and Bioconductor [5] in R, and the scverse project [6] in Python. Despite differences in implementation, the underlying data structures are largely shared across frameworks whereby gene by cell matrices and associated cell metadata are stored within various slots of each object.

**Figure 1.**
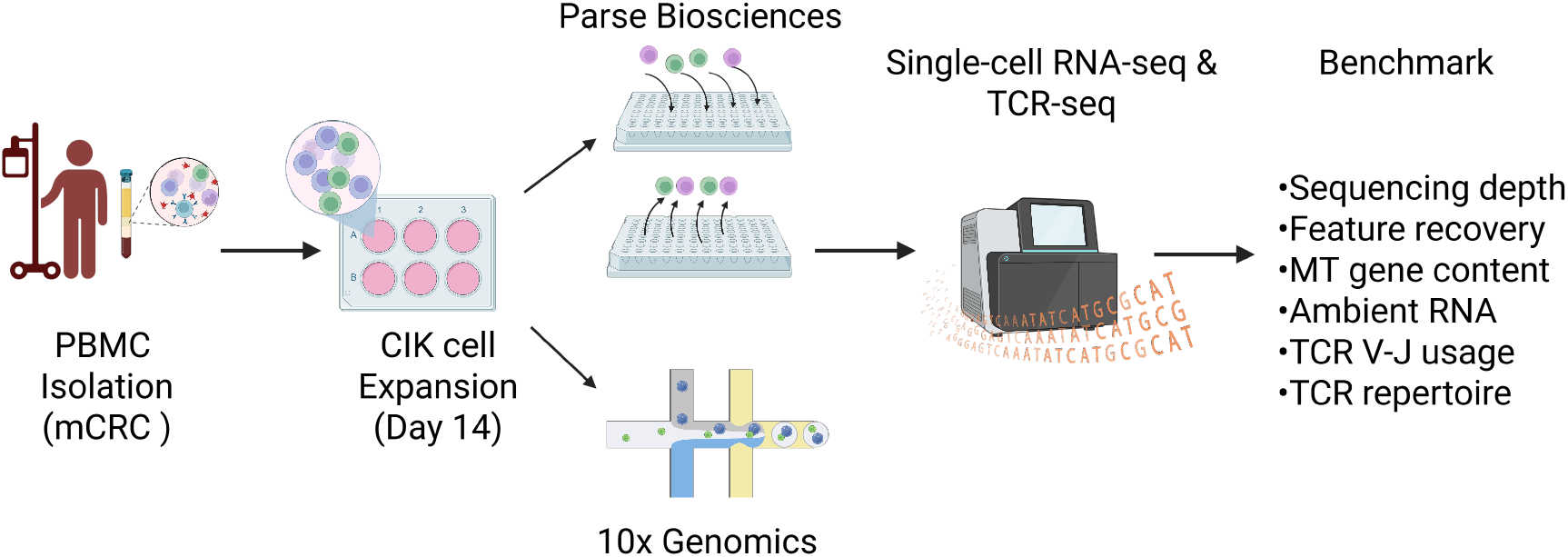
Experimental design and data workflow. CIK cells expanded from PBMC’s from patients with CRC were isolated at day-14 into eight biological and technical replicates. Samples were then processed through 10x and Parse workflows and sequenced. Computational processing, benchmarking and downstream analyses was then performed. *Created with BioRender.com*

This analyses used mainly the Seurat and SingleCellExperiment [7] (SCE) data frameworks due to availability of toolkits for immune profiling analyses and interoperability between object structures.

## 3. Results

### 3.1. Sequencing saturation and downsampling

Sequencing saturation was evaluated by examining how the number of genes detected per cell increases with sequencing depth. Gene detection typically increases with sequencing depth before reaching a plateau, which indicates sequencing saturation. Due to sample pooling and kit specifications, Parse samples achieved approximately 40,000 reads per cell on average across samples, which was approximately twice the sequencing depth of 10x samples.

To reduce systematic bias due to sequencing depth, all Parse samples were downsampled to approximately 20,000 mean reads per cell to match the average sequencing depth of 10x samples across datasets, which can be seen in pipeline summary reports. Samples were down-sampled at the FASTQ level to a target mean of 20,000 reads per cell using the `Seqtk toolkit v1.3` [8]. Targeting mean reads per cell provides an accurate and consistent approach for downsampling single-cell data [9]. Regardless of downsampling, some cells will always return a high count of reads per cell in single-cell experiments. The effects of downsampling is seen on the sequencing depth scatter plot where reads per cell is lower overall (Fig. 2A).

**Figure 2.**
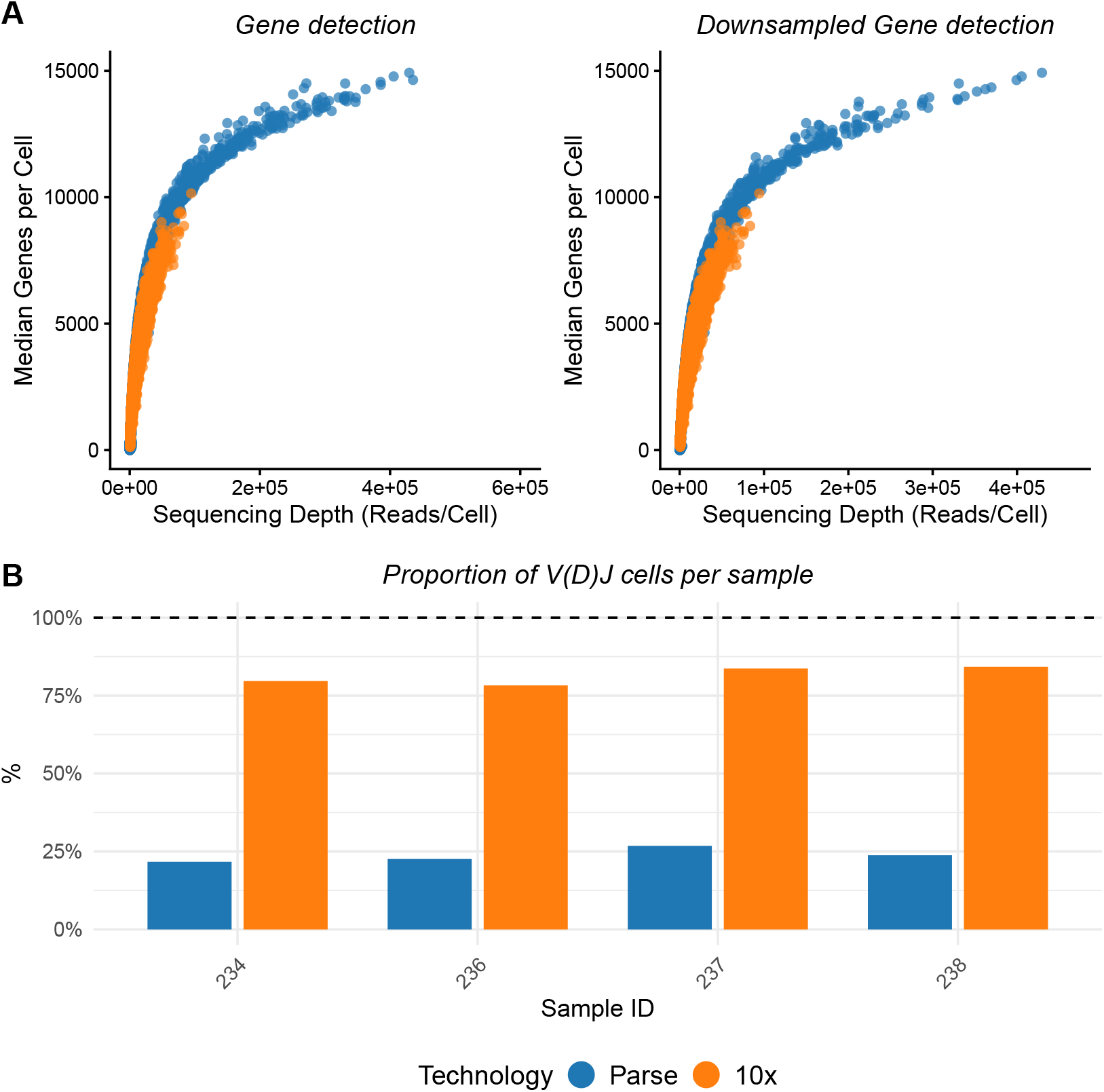
A. Left: Gene detection saturation per technology. Right: Downsampled gene detection per technology. B. Proportion of V(D)J cells per sample.

The recommended sequencing depth for V(D)J libraries is 5,000 reads per cell for both 10x and Parse. Downsampling of single-cell gene expression data does not affect immune profiling results when V(D)J libraries have reached sequencing saturation, as was the case here. Both TCR libraries reached sequencing saturation at the targeted depth of 5,000 reads per cell.

### 3.2. Feature Recovery and QC

To accurately evaluate immune profiling kits, per-cell feature variables should firstly be examined. Vendor pipelines will output per-cell gene and transcript counts. Commonly used QC metrics were calculated including mitochondrial (MT) and ribosomal protein (RP) gene content using the Seurat function `PercentageFeatureSet`. All feature metadata can be conveniently stored in a SCE or Seurat object, allowing for quick and efficient plotting and analysis of feature variables at the sample level. Violin plots per sample will allow for side-by-side comparison of QC metrics. Violin plots often represent commonly used QC metrics and per call variables for single-cell data [7]. Single-cell data objects can be constructed with predefined cut-offs for gene counts per cell but not transcript counts per cell. Cell filtering based on transcript counts would typically be done after object construction which provides the need to define cut-offs during data QC. Algorithms within custom vendor pipelines will also contain pre-defined cut-offs for cell and feature calling.

#### 3.2.1. Transcript and Gene counts per cell

After downsampling, transcript counts per cell are similar across samples. Violin plots of the raw data show that the median remains below 100,000 transcripts per cell for all samples (supplementary figure 1). Parse samples include a smaller subset of cells exceeding 200,000 transcripts per cell, while all 10x samples do not surpass 100,000 transcripts per cell for any sample. Cells with high transcript counts are likely doublets or multiplets and are typically removed from downstream analyses.

Across all Parse samples, with or without downsampling, the data shows a broader distribution at the low end, whereas 10x samples display a sharper cut-off. Rarer transcripts can appear on either end as samples are sequenced deeper. Because single-cell data is inherently sparse, plotting log-transformed counts per-cell provide a clearer view of the end ranges of count distributions (supplementary figure 2).

For gene counts per cell, Parse samples display a broader distribution and include an additional population of cells with lower gene counts whereas 10x samples have a narrower gene count distribution across all samples (Fig. 3A). A similar gene count distribution with outlier cells has been documented elsewhere for Parse data [10]. As seen on the violin plots, all Parse samples exhibit a bimodal distribution, with a distribution nearer zero for all samples that likely represents lower complexity or background noise. In contrast, all 10x samples show a unimodal distribution, characterized by one main density region reflecting consistent gene expression complexity across cells. Downsampling primarily influences the upper range of gene counts per cell in all Parse samples. Following downsampling, Parse samples still show a bimodal distribution and align closer with 10x samples, with mean gene counts per cell converging at approximately 2,500 across all samples.

**Figure 3.**
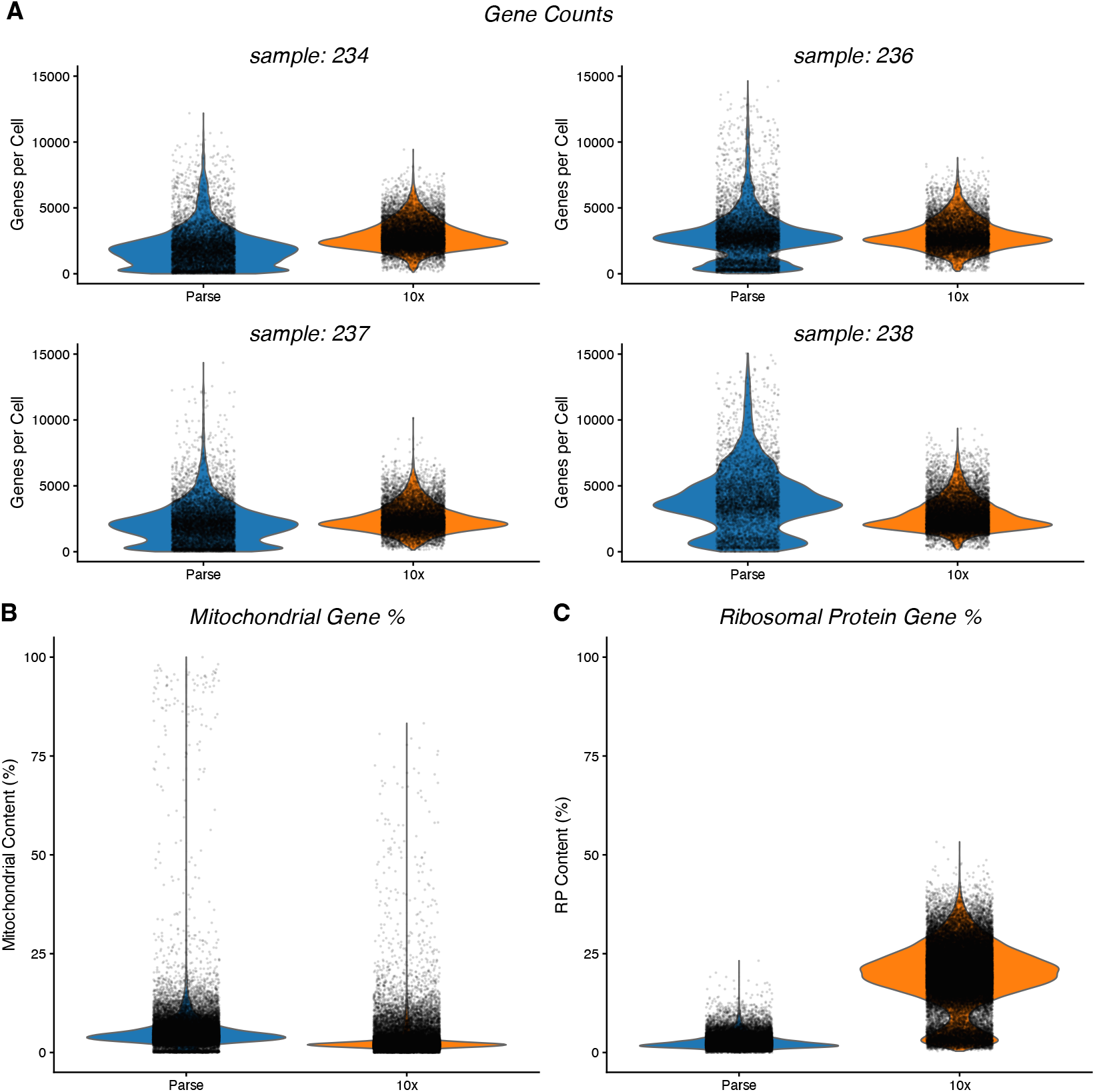
A: Gene counts per cell across sample and technology. B: Mitochondrial gene percentage and C: Ribosomal protein gene percentage content per cell across sample and technology. Parse data is shown in blue, while 10x data is shown in orange.

#### 3.2.2. Highest expressed features

The features with the highest average expression across all cells were plotted for all samples using the `scater` R package and the function `plotHighestExprs()` (supplementary figure 3). This plot can inform which genes contribute the most to the overall expression signal and is useful for identifying dominant transcripts that may influence downstream analyses, especially cell clustering.

Examining these top-expressed genes also helps distinguish between biological signal and technical artifacts, as some genes—such as *MALAT1*, which appeared among the highest expressed in both technologies, can be non-informative housekeeping or artifactual genes [11]. Additional clustering analyses and sample-specific context is required to determine whether certain highly expressed genes represent genuine biology or technical noise. Other commonly detected features in both datasets included *B2M, GAPDH, HLA-A*, and *HLA-B*.

#### 3.2.3 Mitochondrial and Ribosomal Protein content

Mitochondrial (MT) and Ribosomal Protein (RP) gene content per cell was compared between Parse and 10x to evaluate differences in QC metrics. Examining these features is an established QC step in single-cell analyses as high expression indicates lower quality cells or cells that are under stress [12]. MT genes begin with ‘MT-’ in human data and ‘mt-’ in mouse data, whereas RP genes are labeled with ‘RPS’ or ‘RPL’.

Visualization with violin and boxplots showed that while Parse samples have more variation in MT content, most were broadly comparable to 10x samples, with the exception of sample 234, which exhibited noticeably higher mitochondrial levels and contributed substantially to the observed difference. Samples were pooled across technologies and then plotted (Fig 3B,C) to get an overview of the library. Parse data returned cells with higher MT gene counts overall with some cells nearing 100% MT content (supplementary figure 4), similar results have also been seen elswehere [10]. Cells expressing high MT content will typically be removed prior to clustering and downstream analyses [20].

In contrast to MT content, RP content per cell was substantially higher in 10x data; Fig.C 3. Although some variability was observed among individual samples, the higher RP signal in 10x was consistent overall at approximately 40%, suggesting platform-specific differences in RP capture. The observed difference in ribosomal content likely reflects fundamental distinctions in how the two technologies capture and process RNA. 10x, which relies on droplet-based lysis, releases the entire cellular RNA pool into solution, capturing a broad range of cytoplasmic transcripts, including canonical ribosomal protein mRNAs, and typically resulting in a ribosomal fraction of 20–40%. In contrast, Parse uses an in situ combinatorial barcoding approach, where fixation and multiple washing steps during sample preparation preserve nuclear RNA while removing much of the cytoplasmic RNA. This leads to a lower proportion of ribosomal transcripts, often below 10%.

#### 3.2.4 Ambient RNA contamination

In scRNA-seq, ambient RNA contamination arises when free-floating RNA molecules from lysed cells are captured along with true cellular transcripts, leading to misleading gene expression signals. To estimate contamination levels across all samples, the `DecontX`[13] package was used, as it is platform-agnostic and provides a fair comparison between technologies, unlike methods that are tailored for droplet based methods and depend on empty droplets for contamination estimation.

Filtered and unfiltered outputs from vendor pipelines were analyzed using the default parameters with the function `decontX()`. Contamination levels were then visualized using `plotDecontXContamination()`, which maps contamination scores onto UMAP space (supplementary figure 5). A histogram of contamination scores shows that 10x samples have higher contamination scores overall, indicating that 10x data have slightly more ambient RNA contamination overall, although the difference between technologies is modest.

### 3.3 The Immune Repertoire

Cells are classified as T or B cells by vendor pipelines based on the detection of targeted V(D)J transcripts. To improve cell calling, these pipelines filter out barcodes not identified as cells in the corresponding GEX library, thereby minimizing overcalling. Because the GEX library captures all polyadenylated mRNA—including VDJ-T or VDJ-B transcripts selectively amplified to generate the V(D)J library—it provides greater sensitivity for detecting true cells. This integration ensures that only barcodes confirmed as cells in the GEX data are retained in the V(D)J data, requiring both libraries to be sequenced from the same parent sample, as implemented here for both Parse and 10x immune profiling pipelines.

#### 3.3.1 T cell proportions

The proportion of cells identified as V(D)J cells per sample was analysed (Fig. 2B). Both pipelines define V(D)J cells as cell barcodes that express targeted V(D)J transcripts. Evaluating the number and proportion of these calls of T or B cells in each sample is done by extracting relevant clonotype information from respective pipeline outputs. It can be seen that 10x returns a higher proportion of T cells across all samples.

#### 3.3.2 Clonotype distribution

In single-cell immune profiling, a clonotype refers to a unique T or B cell characterized by an identical antigen receptor sequence, defined by the rearranged variable V(D)J gene segments and the resulting complementarity determining region 3 (CDR3). Each respective pipeline identifies the most frequent clonotypes, with the top 10 displayed in the summary report and available in the output files. The top clonotypes per technology were then compared as per pipeline outputs. While the top 10 clonotypes detected were identical across technologies for all samples, the top 50 or 100 clonotypes differed considerably, sharing only about 25% overlap.

Both technologies process matched samples, so it is expected that the most abundant clonotypes appear in both datasets. However, the overlap is not exact, as each technology may detect unique clonotypes depending on differences in sensitivity, sequencing depth, or clonotype-calling methods and this can be expected for most immune profiling technologies.

#### 3.3.3. Clonal length distribution

Clonal length refers to the distribution of CDR3 sequence lengths in single-cell immune receptor data, typically measured in amino acids (aa) or nucleotides (nt). This distribution can be visualized using the `clonalLength()` function from the `scRepertoire`[16] package.

The observed lengths often display a multimodal pattern, reflecting differences in the number of TCR chains recovered per cell. For example, a peak around 12 amino acids usually represents cells where only one chain (commonly TCRα or TCRβ) was recovered. This can be due to incomplete capture during library preparation, sequencing inefficiencies, or true biological states such as immature T cells expressing only a single chain [14]. A second prominent peak between 25–30 amino acids typically corresponds to conventional T cells in which both TCRα and TCRβ chains were successfully captured, with their combined CDR3 lengths summing to that range. A third component, a tail extending above 35 aa suggests the presence of multiple TCR chains within a single-cell barcode. This can arise from biological phenomena such as dual TCRα (common) or dual TCRβ (less common) [15], or from technical artifacts like doublets or barcode collisions. For example, a sequence such as CALSTNFGNEKLTF;CAA SGTGGYKLTF_CASSRGTGSYEQYF;CASSPAGDQ PQHF includes multiple alpha and beta chains, with the underscore separating the TCRα (left) and TCRβ (right) components. Such extended CDR3 lengths, and the structure of these combined sequences, offer valuable insight into clonotype diversity, pairing fidelity, and overall data quality.

A clonal length histogram was plotted to compare CDR3 sequence lengths between all samples (Fig. 4A). The histogram revealed that 10x data consistently showed higher distributions across the length range. In particular, a clear peak around 25–30 aa was more prominent in the 10x data, suggesting a greater number of cells with recovered conventional paired alpha-beta TCR’s. A secondary peak above 35 aa was also more evident in 10x data, indicating the presence of more multichain cells or more than two contigs per cell, which could reflect true dual TCR expression or potential doublets. In contrast, both technologies showed similar distributions at around 12 aa peak, consistent with cells where only a single chain was recovered.

To investigate TCR chain bias, the `clonalLength()` function was used to plot specific TCR chains by using the `chain` argument. TCRα has a lower expression than TCRβ across both technologies; Fig. 4B. This observation is important for researchers conducting chain-specific analyses, for example in studies examining TCRα-driven responses to COVID-19 vaccination[17].

**Figure 4.**
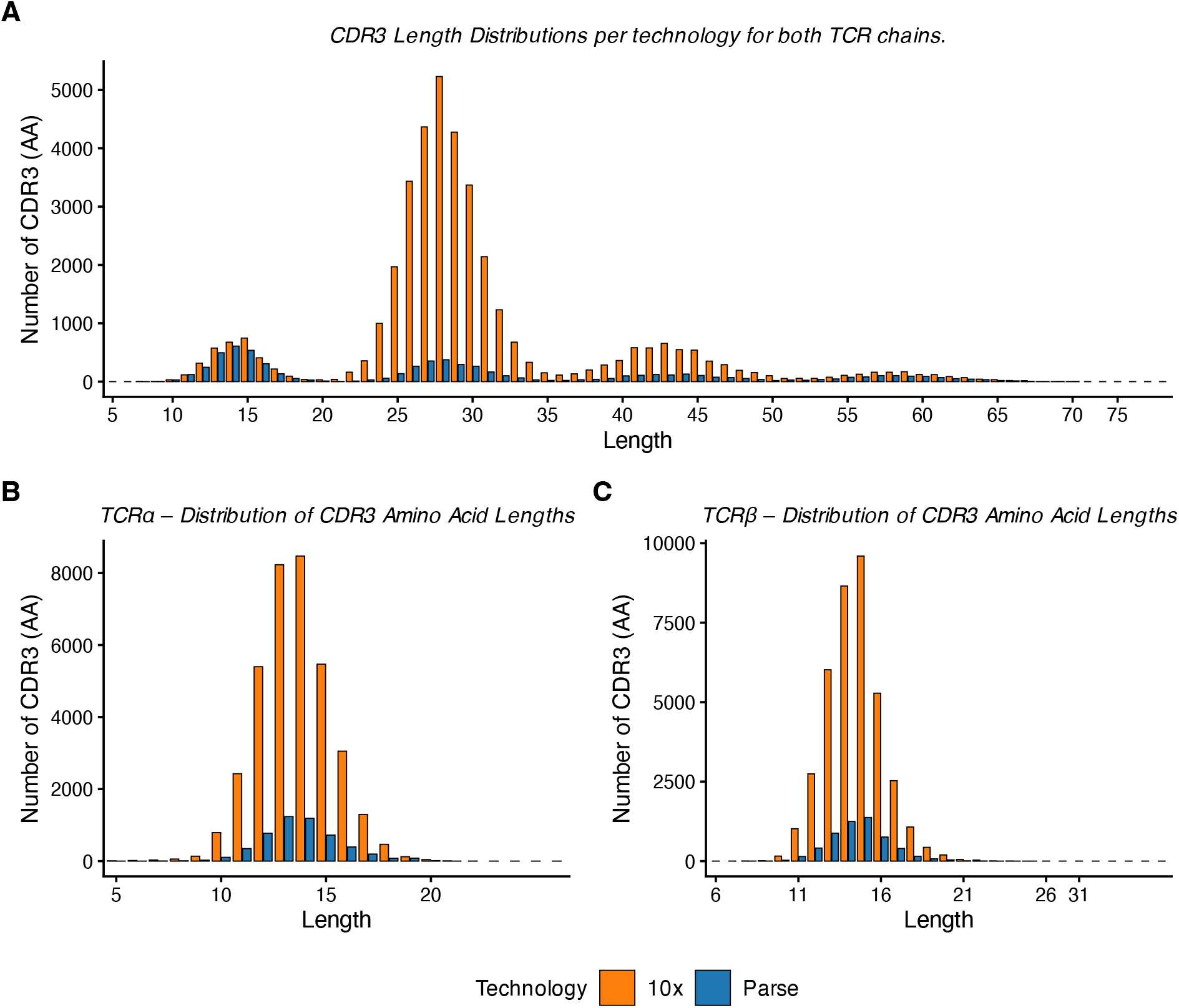
Distribution of CDR3 amino acids across technologies. Top: Both TCR*α* and TCR*β* chains plotted showing a larger distribution of conventional paired alpha-beta TCR’s at ∼ 25aa for 10x Genomics. The smaller population at ∼ 12aa is single-chain cells which shows a more comparable distribution across technologies. The larger distribution indicates the recovery of more than two contigs per cell. Bottom left: TCR*α*. Bottom right: TCR*β*.

#### 3.3.4. TCR V–J Gene Usage Across Samples

V–J gene segment composition was analyzed to compare TCR gene usage across samples and technologies. The `percentVJ()` function, again from the `scRepertoire` package, can calculate, for each sample, the proportion of V and J gene segment usage within a given chain (first α, then β). This generates a numeric table where each row represents a sample and each column represents the relative usage of specific V–J gene combinations. A principal component analysis (PCA) is then performed on this table to summarize the main sources of variation in V–J gene usage patterns across samples. Finally, samples are plotted on PCA space, where samples positioned closer together share more similar V–J gene usage profiles. Separate PCA plots are created for the TCRα and TCRβ chains to compare how the TCR repertoire composition differs between samples and technologies.

The PCA plot of TCR V–J gene usage revealed distinct grouping patterns between the two chains. For the TCRα chain, samples clustered primarily by technology, indicating that technical factors such as capture efficiency strongly influences TCRα V–J profiles Fig. 5. In contrast, the TCRβ PCA showed samples clustering in their matched biological sample, suggesting that TCRβ V–J usage is more consistent within biological replicates and less affected by platform-specific biases. These results indicate that technical bias affects TCRα more strongly, while TCRβ better reflects true biological similarity.

**Figure 5.**
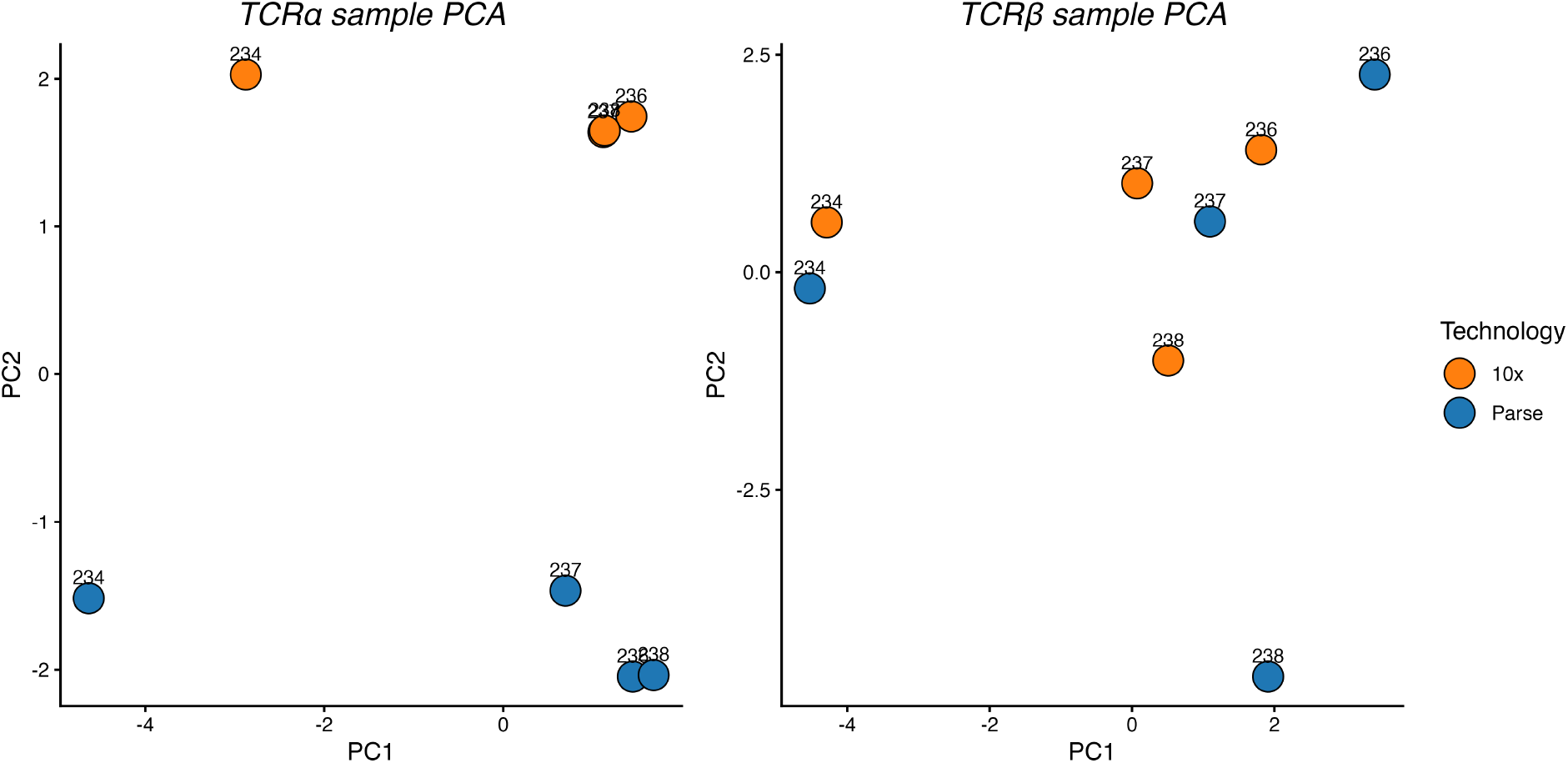
Principal component analysis of TCR V–J gene usage across samples and technologies. Left: TCR*α* chains, where samples group by technology. Right: TCR*β* chains, where samples group by biologically matched pairs.

## 4. Discussion

Benchmarking single-cell immune profiling technologies is essential to understand their performance characteristics and to guide accurate data interpretation and experimental design. In this comparative analysis of 10x Genomics and Parse Evercode, both platforms generated high-quality immune receptor and gene expression data; however, clear and consistent differences were observed in capture efficiency, RNA composition, and chain-specific recovery. Sequencing saturation was reached for both V(D)J libraries, indicating that sequencing depth did not influence clonotype detection while gene expression libraries required downsampling to achieve comparability between technologies. These findings also underscore the importance of sufficient sequencing depth for single-cell gene expression libraries. Although V(D)J libraries reached saturation at recommended depths, deeper sequencing of gene expression libraries on the order of 40,000–50,000 reads per cell is preferable to capture the majority of expressed features.

Despite these adjustments, systematic differences remained. 10x datasets consistently showed higher ribosomal protein (RP) content in compsrison to Parse data for all samples. Ambient RNA contamination was slightly higher in 10x samples, although the overall difference between technologies was modest. Parse data exhibited a broader dynamic range in both reads and transcripts per cell, particularly at the high and low extremes. However these counts would likely be filtered during QC steps of downstream bioinformatic analyses. Rarer transcripts were more apparent at deeper sequencing depths, but gene counts did not double in Parse samples despite increased sequencing, suggesting diminishing returns at higher depth. The broader distribution of transcript counts, especially in violin plots, makes defining QC cut-offs more challenging for Parse data. While log-transformed visualizations help clarify these distributions, some findings may also reflect differences in vendor computational processing pipelines and predefined thresholds, which can influence downstream analyses. However these are the available pipelines to the majority of end users. A common pipeline to process all data was not available for single-cell immune profiling at this time. Using a common pipeline could yield more comparable results and would be useful for analysts that are handling samples across technologies.

In terms of immune repertoire recovery, 10x captured more TCR sequences overall compared to Parse across all samples. Both technologies recover a comparable number of single-chain cells (TCRα or TCRβ), whereas 10x clearly returns much more conventional dual TCRα-β cells and multi-chain cells and this has been shown elsewhere [20]. A chain-specific bias was observed in both technologies, with TCRα consistently detected at lower levels than TCRβ and this has been shown in bulk approaches to TCR capture [18]. This bias was further reflected in V–J gene usage analyses: TCRα samples clustered primarily by technology, indicating stronger technical influence, whereas TCRβ samples grouped by biological replicate, suggesting closer representation of underlying biology. This relates to the importance of establishing a reliable ground truth and reflects a common limitation in computational benchmarking, as has been shown elsewhere for V(D)J repertoire analyses [19]. Despite these differences, strong agreement was found in the most abundant clonotypes, with the top ten clonotypes largely shared between technologies, while overlap decreased among less frequent clonotypes.

Another limitation of this comparison is the differing sample handling requirements between Parse and 10x. Parse relies on fixation, enabling useful sample preservation and batching but potentially introducing RNA degradation or accessibility biases. In contrast, 10x requires fresh cells, better reflecting native transcriptional states but increasing susceptibility to dissociation stress and handling variability. These pre-analytical differences can impact gene detection, cell type representation, and the immune repertoire, complicating direct platform comparisons.

Together, these findings highlight the importance of understanding platform-specific characteristics and clear research questions when designing, analysing and interpreting single-cell immune profiling experiments. Systematic technological biases, similar to those reported in bulk sequencing comparisons, can influence gene expression metrics, chain recovery, and repertoire composition. Careful benchmarking and transparent reporting of parameters such as sequencing depth, chain recovery rates, and contamination levels are essential to ensure reliable and comparable results across technologies.

### 4.1. Data

Data analyses: https://github.com/cathalgking/single-cell-immune-repertoire-benchmarking

## 4.2. Acknowledgements

We would like to thank the vendors for their technical support and the South Australian Genomics Centre (SAGC) for their services. Funding was provided to support the single cell sequencing work by Cure Cancer Australia (K.F) and a Cancer Council SA Mid-Career Research Fellowship (K.F). We would also like to thank the clinical staff and patients from The Queen Elizabeth Hospital who supported this work.

## Notes

### Competing Interest Statement

The authors have declared no competing interest.

