## Supplemental Figures for "Comparative benchmarking of single-cell transcriptomes and immune repertoires across technologies"

Supplementary figures

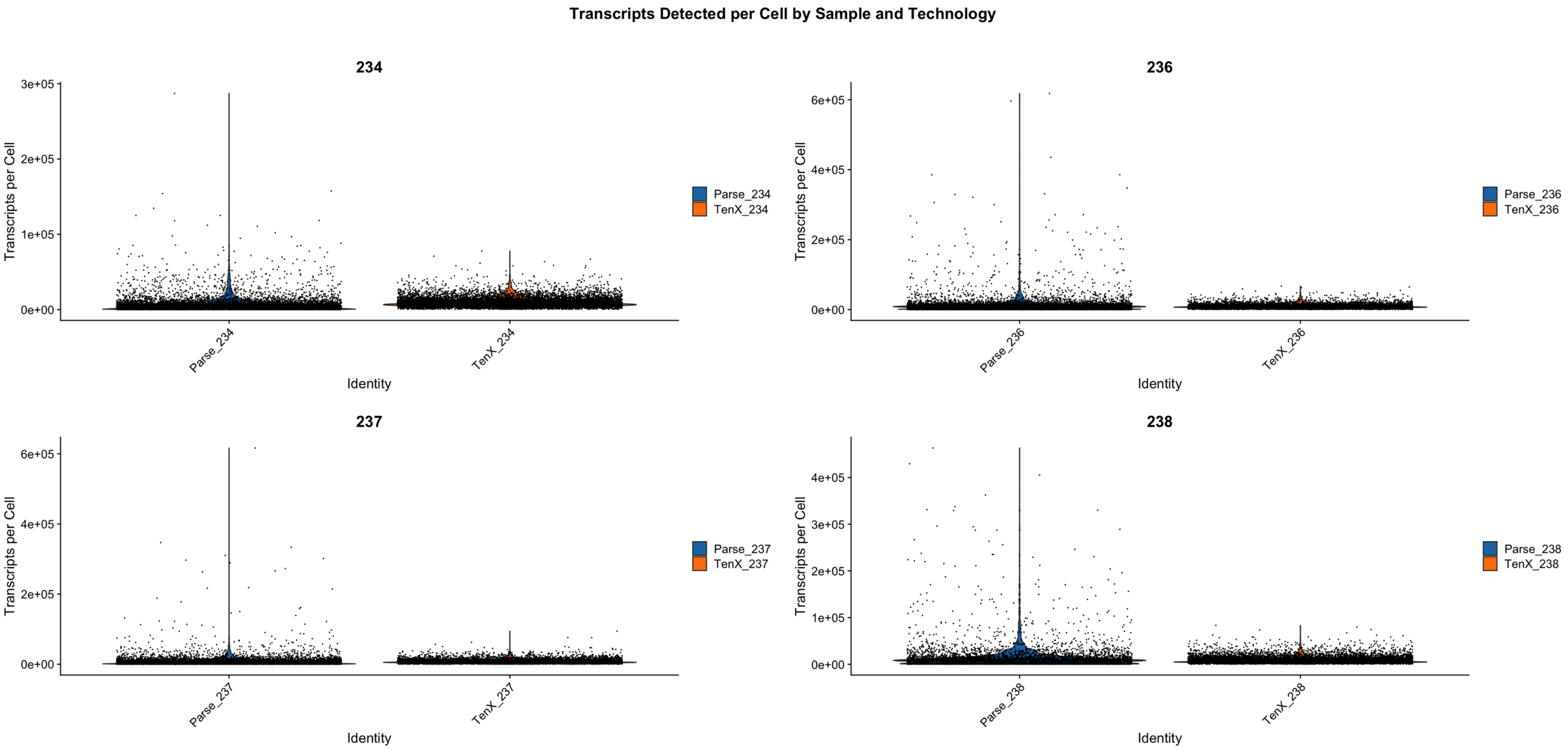


Figure 1: Transcript counts per cell by sample and technology.

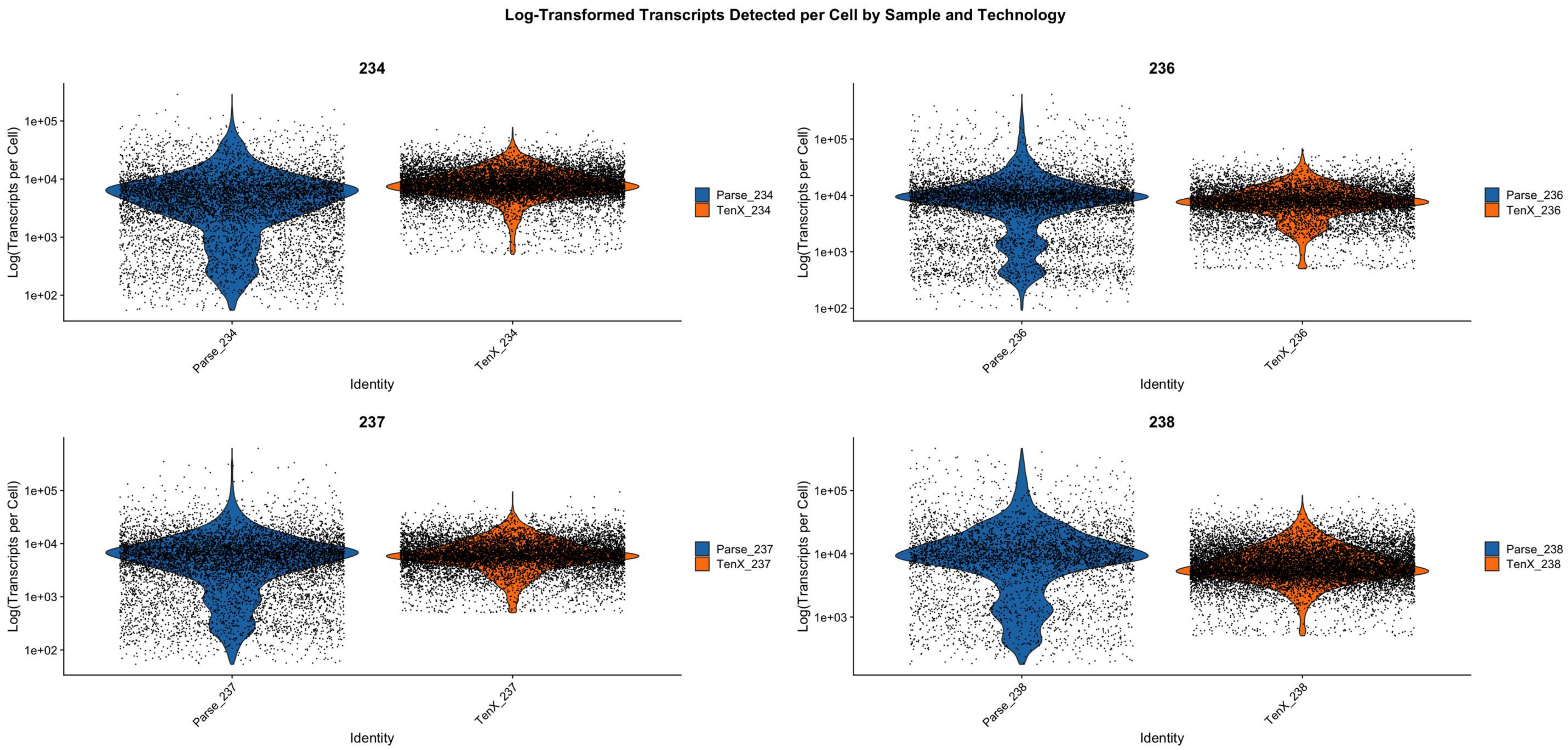


Figure 2: Log transformed transcript counts per cell by sample and technology.

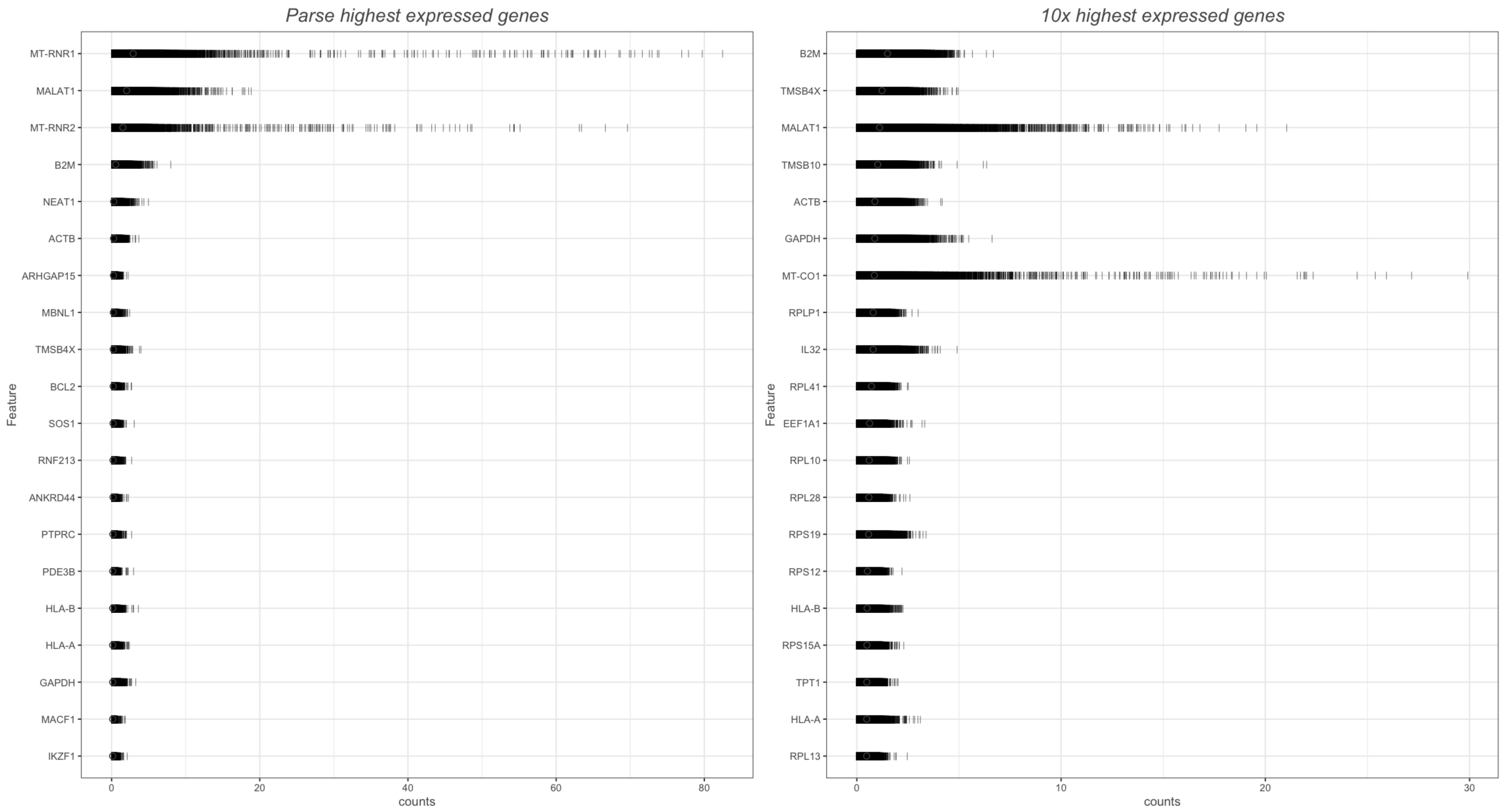


Figure 3: Highest average expressing genes. All samples pooled per technology with Parse on the left and 10x on the right.

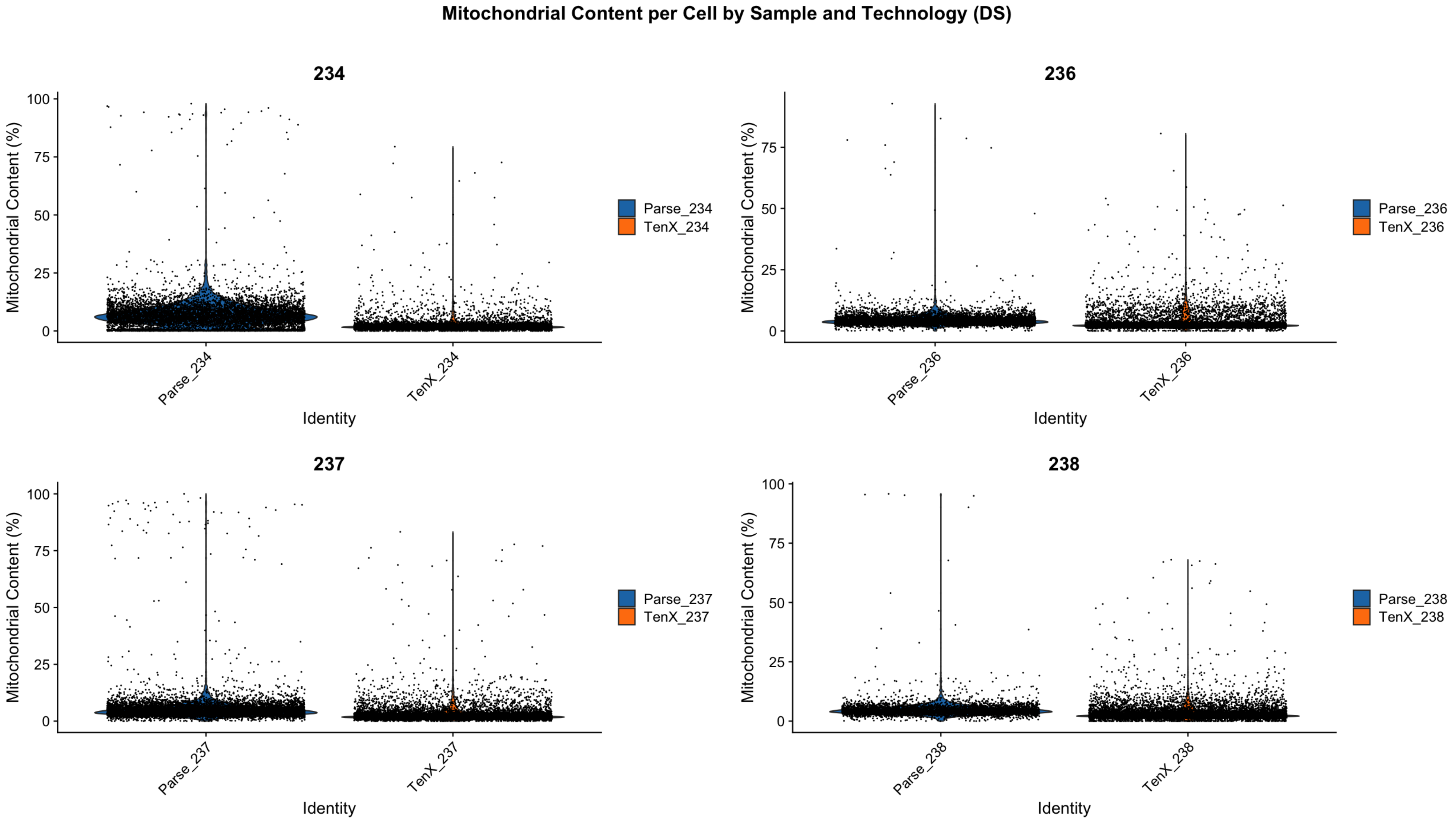


Figure 4: Mitochondrial counts per cell by sample and technology.

### Ambient RNA contamination

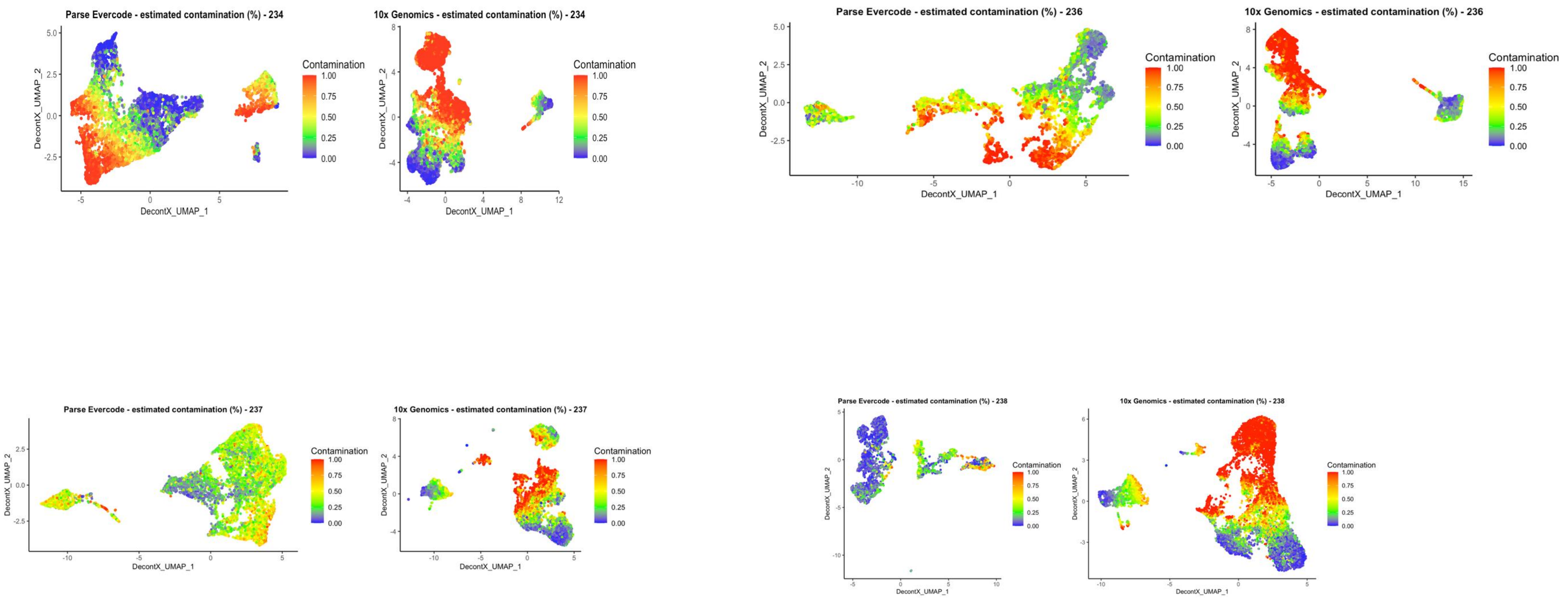


Figure 5: Estimated Ambient RNA contamination levels by sample and technology.
